# *Wasf1* deletion attenuates tau hyperphosphorylation, microglial state transition, and cognitive deficits in P301S tau mice

**DOI:** 10.64898/2026.01.14.699391

**Authors:** Clara Berdasco, Woo-Hyun Cho, Jordan Lee, Annu Annu, Jose H. Ledo, Yu Young Jeong, Yoon-Seong Kim, Steven S. An, Yoshiaki Tanaka, Jung-Hyuck Ahn, Yong Kim

## Abstract

We previously reported that WAVE1, a major activator of Arp2/3 complex-mediated actin polymerization, is downregulated in postmortem brains of Alzheimer’s disease (AD) and that WAVE1 regulates amyloid precursor protein trafficking and amyloid-β production. However, its role in tau pathology remains unknown. Here, we demonstrate that WAVE1 activity is suppressed in P301S tau mice through elevated inhibitory phosphorylation. Strikingly, WAVE1 gene (*Wasf1*) knockout in P301S tau mice significantly reduces tau hyperphosphorylation and improves cognition, suggesting a compensatory role for WAVE1 suppression in tau pathogenesis. Single-nucleus RNA sequencing reveals that *Wasf1* deletion in P301S tau mice reverses microglial state transitions, with minimal impact on other brain cell types. *Wasf1* mRNA is highly translated in microglia in non-Tg mice, while its expression is downregulated in P301S tau mice. *Wasf1* knockdown in BV2 microglia cells enhances the degradation of engulfed tau fibrils, indicating WAVE1 as an endogenous regulator of microglial function. Additionally, CellChat analysis indicates that *Wasf1* deletion alters microglial autocrine signaling and their interactions with other cell types in P301S tau mice. These findings, taken together, suggest that *Wasf1* deletion restores homeostatic microglial function, mitigates tau pathology, and alleviates cognitive deficits, highlighting WAVE1 as a potential therapeutic target for tauopathy-related dementias.

## Introduction

Actin polymerization and depolymerization are fundamental processes that govern functional adaptation in neurons and neuroglia, including microglia, in the brain. Actin dynamics are tightly regulated by numerous actin-binding proteins, including WAVE regulatory complex (WRC), a heteropentameric complex composed of WAVE, NCKAP, CYFIP, ABI, and BRK1 (1). As the core functional subunit, WAVE activates actin-related protein 2/3 (Arp2/3) complex-mediated actin polymerization while the other subunits modulate WRC assembly, localization, and upstream signaling (2–5). Of note, WAVE1 (Wiskott-Aldrich syndrome protein family verprolin-homologous protein 1) is a brain-enriched paralogous protein derived from the ancestral WAVE gene and plays critical roles in synaptic development and plasticity, as well as regulation of mitochondrial morphology and trafficking in neurons (6, 7). In addition, WAVE1-mediated actin polymerization is essential for oligodendrocyte morphogenesis and axonal myelination (8). However, the role of WAVE1 in microglia, particularly under disease-associated conditions, remains largely unexplored.

Previously, we identified the WAVE1-containing WRC as a binding partner of cyclin-dependent kinase 5 (Cdk5) and its regulatory subunit, p35, in brain tissue, and established WAVE1 as a substrate of Cdk5/p35 (9). Cdk5, also known as a tau kinase (10), is dysregulated in Alzheimer’s disease (AD) (11). Cdk5 phosphorylates three sites on WAVE1 (Ser310, Ser397, and Ser441), and WAVE1 phosphorylation in these sites inhibits WAVE1-mediated activation of Arp2/3-dependent actin polymerization in *in vitro* assays (9). In cells and brain tissue, the baseline stoichiometry of Cdk5 phosphorylation on WAVE1 is high, leading to the inhibition of WAVE1 activity. Upon stimulation of dopamine D1 receptors or NMDA glutamate receptors, WAVE1 is dephosphorylated at all three sites, thereby becoming activated through cAMP and Ca²⁺-mediated activation of protein phosphatase-2A and protein phosphatase-2B, suggesting WAVE1 as a neuronal activity-dependent regulator of actin polymerization (9, 12).

Alterations of the WRC have been implicated in neurodegenerative diseases, including AD and Parkinson’s disease (1). WAVE1 has emerged as a key AD-associated protein in both human postmortem brains (13) and mouse models (14). Our previous work demonstrated consistent downregulation of WAVE1 across human AD brains, as well as in AD mouse and cell models (15). Mechanistically, the amyloid precursor protein (APP) intracellular domain (AICD), generated via the amyloidogenic pathway, binds to the WAVE1 gene (*WASF1*) promoter and suppresses its transcription and protein expression (15). Functionally, WAVE1 interacts with APP in the Golgi apparatus and regulates vesicle budding and trafficking, thereby increasing Aβ production (15). *Wasf1* knockout (KO) reduces Aβ levels and rescues cognitive deficits in APP/PS1 mice, indicating a protective effect of WAVE1 reduction (15). However, whether WAVE1 is altered in the context of tau pathogenesis and whether it contributes to the progression of tau pathology remain unknown.

Notably, a coding variant in *ABI3*, a microglia-enriched paralog of the ABI subunit in the WRC, has been identified as a genetic risk factor for late-onset AD (16, 17). *Abi3* KO impairs microglial motility and phagocytosis, exacerbating Aβ and tau pathology in AD models (18–20). Interestingly, WAVE1 has also been shown to regulate innate immune responses in peripheral macrophages (21). However, the role of WAVE1 in the functional state of microglia, the brain’s resident macrophages, and its relationship to AD pathogenesis remain largely unexplored. In this study, we uncovered a beneficial role for WAVE1 deletion in the P301S tauopathy model. *Wasf1* KO attenuates tau hyperphosphorylation, improves cognition, and reverses the disease-associated microglia (DAM) gene signature in P301S tauopathy mice. These microglial changes may be driven by both cell-autonomous and non-cell-autonomous mechanisms. Taken together, our findings suggest that WAVE1 represents a promising therapeutic target across AD pathologies.

## Results

### Wasf1 KO reduces tau hyperphosphorylation and gliosis in P301S tau mice

To investigate the role of WAVE1 in tau pathology, we used an Aβ-independent tauopathy model, given WAVE1’s critical role in APP trafficking and Aβ production (15). The P301S mutation in tau is associated with familial frontotemporal dementia (22) and promotes tau detachment from microtubules, facilitating fibril formation (23). We employed P301S tau mice (PS19 model) (24) and examined WAVE1 changes using immunoblotting. WAVE1 protein levels were unchanged. However, WAVE1 phosphorylation was significantly increased in both female and male P301S tau mice compared to respective non-transgenic (non-Tg) controls (Figure 1, A-C), suggesting suppression of WAVE1 activity and WAVE1-mediated actin polymerization via inhibitory phosphorylation (9). To assess the impact of WAVE1 suppression on tau hyperphosphorylation, we crossed P301S tau mice with constitutive *Wasf1* KO mice (15). *Wasf1* deletion significantly reduced tau hyperphosphorylation at AD-relevant sites S202/T205 (AT8) and T231 (AT180), accompanied by increased total tau levels (Figure 1, D-G). Phosphorylation at S199 was not significantly affected. The results were consistent in both sexes. Previous studies indicate microglial activation plays a critical role in tau pathogenesis in P301S tau mice (24, 25). Consistent with this, we observed an increase in microglial numbers (Iba1-positive cells) in P301S tau mice compared to non-Tg controls. *Wasf1* KO reversed microgliosis in both males and females (Figure 1, H-K).

**Figure 1.**
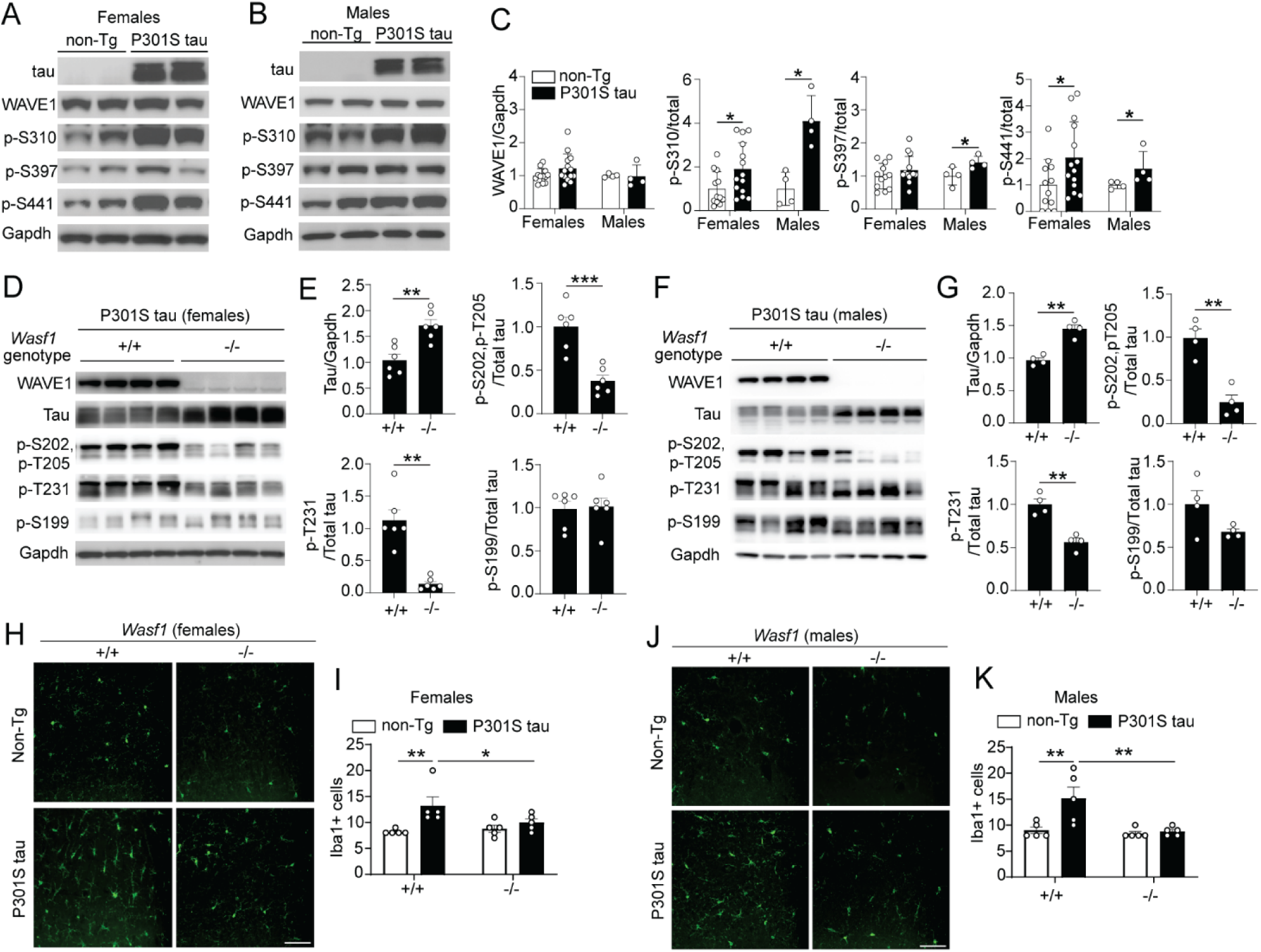
WAVE1 is hyperphosphorylated in P301S tau mice, and *Wasf1* KO mitigates tau hyperphosphorylation and gliosis in P301S tau mice. (**A-C**) The levels of tau, WAVE1, p-WAVE1 (p-S310, p-S397, p-S441) and Gapdh in hippocampal lysates from non-Tg and P301S tau mice were analyzed by immunoblotting. Representative blot images of females (**A**) and males (**B**), and the quantification results of females (n=12-14/group) and males (n=4/group) (**C**). (**D**-**G**) Levels of p-tau (AT-8: p-S202/p-T205; AT-180: p-T231; p-S199), total tau and Gapdh in hippocampal lysates from P301S tau mice harboring *Wasf1* WT (+/+) or KO (-/-) were analyzed by immunoblotting. Representative blot images of females (**D**) and males (**F**), and the quantification results of females (**E**, n=5-6/group) and males (**G,** n=4/group). (**H**-**K**) Representative images of Iba1 in females (**H**) and males (**J**). Scale bars, 100µm. The number of Iba1-positive cells in CA1 from non-Tg and P301S tau mice harboring *Wasf1* +/+ or -/- alleles was quantified. (**I**, females, and **K**, males; n=5 mice/group). Females and males aged 8-9 months were used for all experiments. Data are presented as means ± SEM. \**p*<0.05, \*\**p*<0.01, \*\*\**p*<0.001, t-test (C, E, and G), and two-way ANOVA Tukey’s *post hoc* test (I and K).

### Wasf1 KO improves cognition in P301S tau mice

We next examined the effects of *Wasf1* KO on cognitive behavior. Because homozygous *Wasf1* KO causes sensorimotor deficits (26), we evaluated heterozygous *Wasf1* KO mice to avoid confounding effects on behavioral outcomes. Locomotor activity in non-Tg control mice and P301S tau mice with WT *Wasf1* or heterozygous *Wasf1* KO was comparable in the open-field test (Figure 2, A and B; Supplementary Figure 1, A-D). P301S tau females (Figure 2C), but not males (Figure 2D), exhibited cognitive deficits compared to non-Tg controls in the Y-maze test (working memory). *Wasf1* heterozygous KO significantly ameliorated these deficits in P301S tau females (Figure 2C). In the Morris water maze (spatial memory) test, training for hidden platform acquisition was comparable across all groups, indicating intact learning ability (Figure 2, E and G). However, significant memory deficits, measured by swimming time in the target quadrant, were observed in both male and female P301S tau mice compared to non-Tg controls. Of note, *Wasf1* KO significantly improved performance in P301S tau mice (Figure 2, F and H). Locomotor activity during the memory test was comparable across groups (Supplementary Figure 1, E and F). Altogether, these results indicate that *Wasf1* KO confers beneficial effects on cognitive deficits in P301S tau mice.

**Figure 2.**
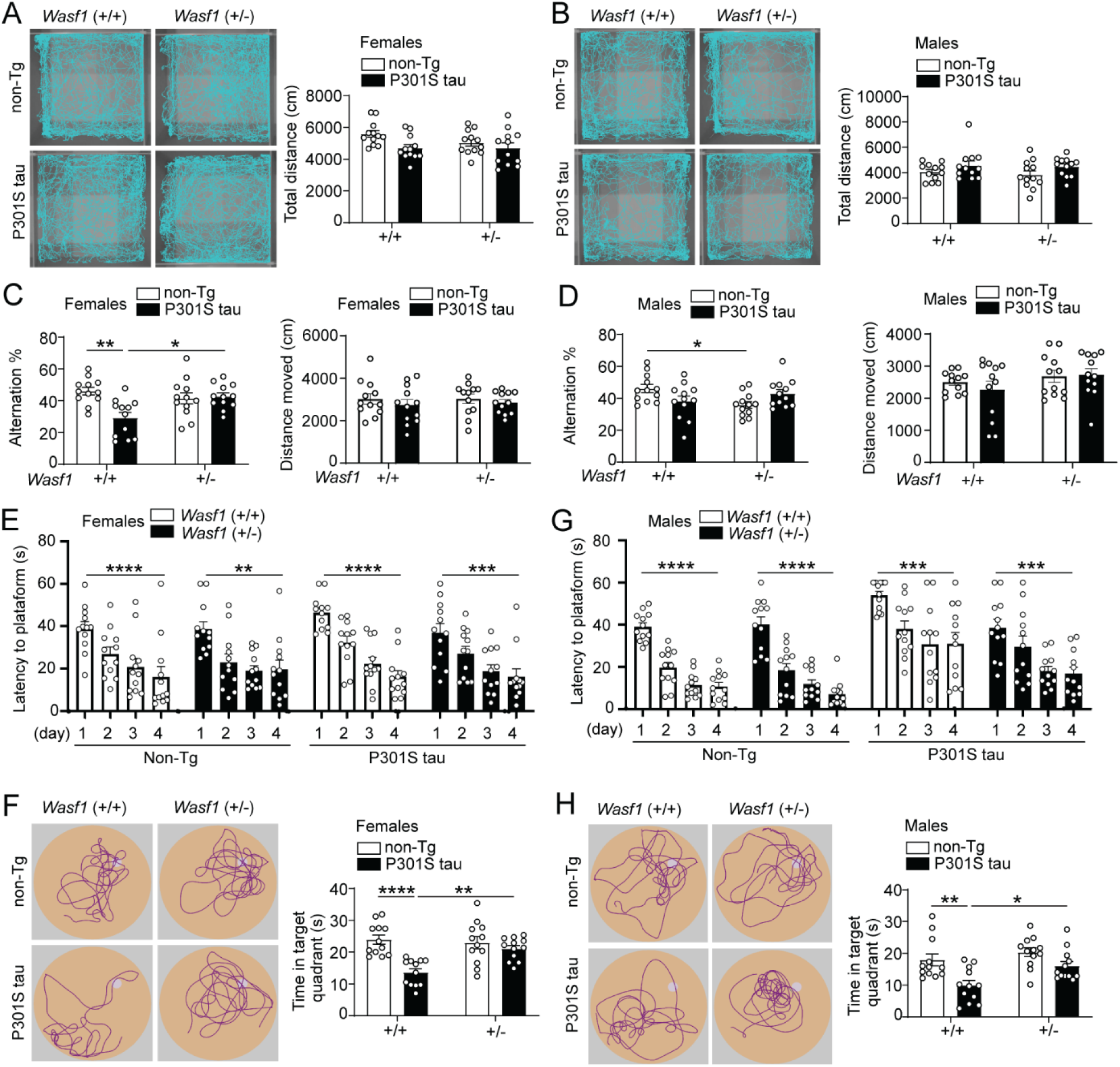
*Wasf1* heterozygous KO improves cognitive performance in P301S tau mice. (**A** and **B**) Effects of WAVE1 reduction on locomotor activity in females (**A**) and males (**B**). Video tracking was recorded (left), and total distance moved (right) was quantified. (**C** and **D**) Spontaneous alteration (left) and total distance moved (right) in the Y maze test for females (**C**) and males (**D**). (**E**-**H**) Morris water maze test showing comparable learning between groups in females (**E**) and males (**G**). *Wasf1* heterozygous KO ameliorates memory deficits in P301S tau mice in both females (**F**) and males (**H**). Representative of swim paths during the probe trial (left), and time spent in the target quadrant (right) are shown. Females and males aged 6-8 months (n=12 mice/group). \**p*<0.05, \*\**p*<0.01, \*\*\**p*<0.001, \*\*\*\**p*<0.0001, paired t-test between day 1 and day 4 (**E** and **G**), and two-way ANOVA with Tukey’s *post hoc* test (**F** and **H**).

### Wasf1 KO reverses microglial transition in P301S tau mice

To investigate the mechanisms underlying the beneficial effects of *Wasf1* KO in P301S tau mice, we performed single-nucleus RNA sequencing (snRNA-seq) on hippocampal tissue. Clustering analysis identified excitatory and inhibitory neurons, microglia, astrocytes, oligodendrocytes, and vascular pericytes (Figure 3A). We first examined cell clusters for changes in their proportions in P301S tau mice compared to non-Tg controls, and whether these changes are rescued in *Wasf1* KO (Figure 3B). The proportion of microglia was significantly increased in P301S tau mice compared to non-Tg mice, and *Wasf1* KO reversed this increase (Figure 3C), consistent with immunostaining results (Figure 1, H-K). Microglial subpopulation profiles were markedly altered in P301S tau mice compared to non-Tg controls and were normalized by *Wasf1* KO, making them comparable to non-Tg mice with WT *Wasf1* (Figure 3D). Differential gene expression (DEG) analysis identified 35 upregulated and 4 downregulated microglial genes in P301S tau mice (WT *Wasf1*) compared to non-Tg controls (WT *Wasf1*) (Figure 3E and Supplementary Table 1). Notably, *Wasf1* KO reversed 11 of the upregulated genes and additionally upregulated 3 others (Figure 3E), indicating substantial restoration of microglial gene expression profiles. P301S tau mice exhibited a strong disease-associated microglia (DAM) signature, which is a phenotype conserved between mice and humans (27, 28). DAM differentiation occurs in two sequential stages (27, 28), both of which were markedly reversed by *Wasf1* KO (Figure 3, F and G), revealing a previously unrecognized role for WAVE1 in regulating AD-relevant microglial states.

**Figure 3.**
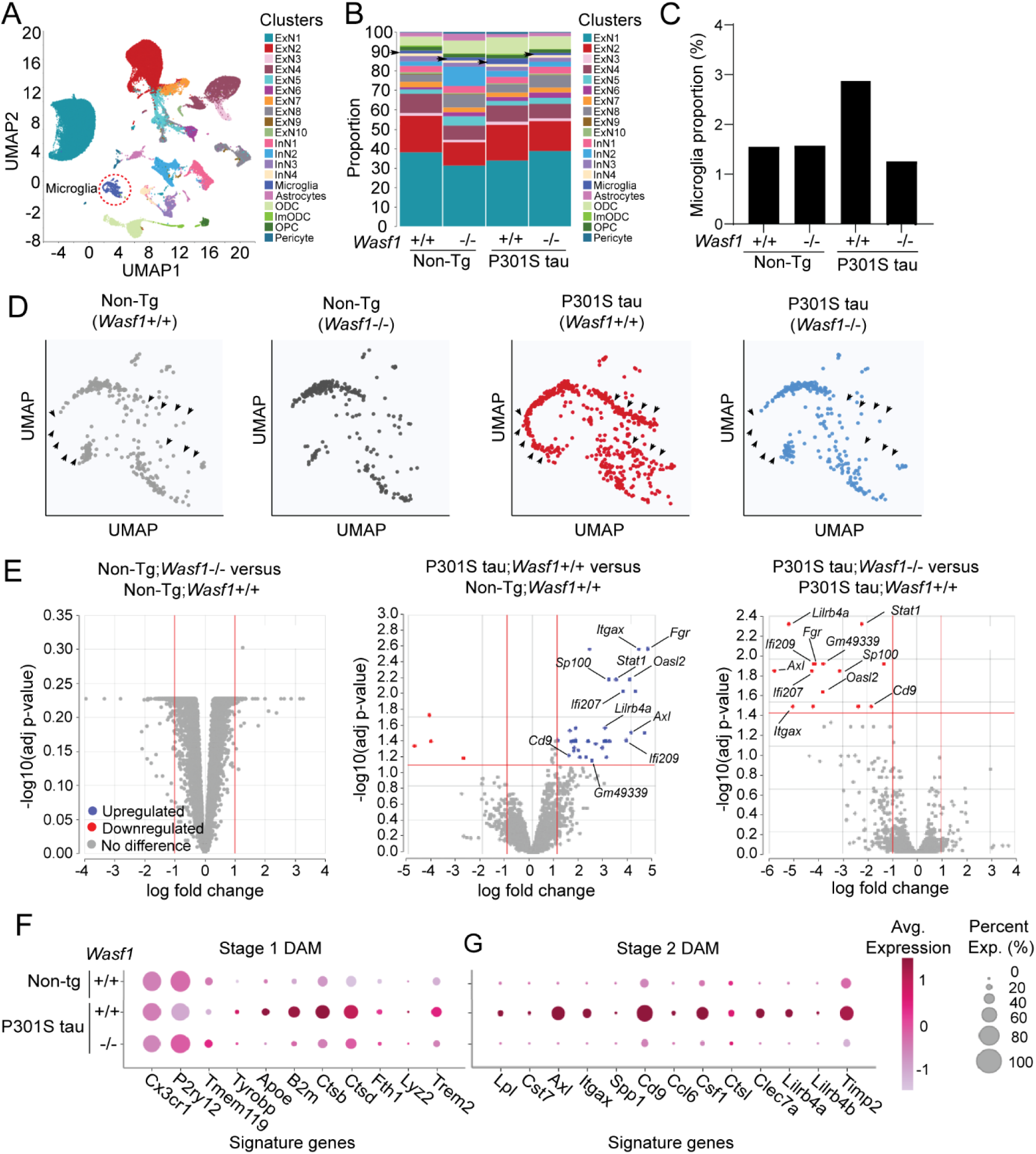
*Wasf1* deletion reverses microglial transitions in P301S tau mice. snRNA-seq was performed on hippocampal samples from non-Tg mice and P301S tau mice with *Wasf1* WT or KO alleles (n = 6/group; 8–9 months old; 3 males and 3 females). (**A**) UMAP visualization of all cell clusters. (**B**) Proportion of specific cell types in each group. (**C**) Proportion of microglia in each group (**D**) UMAP plots highlighting microglial subpopulations. Arrows indicate significant cluster shifts between groups. (**E**) Volcano plots showing DEGs in microglia. The same genes inversely altered by *Wasf1*-/- compared to *Wasf1*+/+ in P301S tau mice are indicated. (**F** and **G**) Expression of representative stage 1 (**F**) and stage 2 (**G**) DAM signature genes.

We have found proportion changes in other cell clusters (e.g., excitatory and inhibitory neurons, oligodendrocytes and astrocytes) (Supplementary Figure 2, C, D, H, I, M, N, R and S) and changes in numerous DEGs between non-Tg mice with *Wasf1* KO and non-Tg mice with WT *Wasf1* (Supplementary Figure 2, E, J, O and T, and Supplementary Table 1). Strikingly, however, a few or no DEGs were detected in these cell types in P301S tau mice compared with non-Tg control mice (Supplementary Figure 2, F, K, P, and U) or *Wasf1* KO versus WT condition in P301S tau mice (Supplementary Figure 2, G, L, Q, and V). These results collectively imply that other cell types might play relatively smaller roles than microglia in mediating the beneficial effects of WAVE1 KO in P301S tau mice.

### WAVE1 is an endogenous regulator of microglia

To investigate WAVE1 protein expression in microglia, we employed a brain immune cell-specific Translating Ribosome Affinity Purification (TRAP) combined with RNA-seq (29), using an immune cell–targeted TRAP line (Figure 4A). Cfh-EGFP-L10a mice express an EGFP-fused large ribosomal subunit (L10a) under the control of the *Cfh* (Complement Factor H) promoter. We isolated ribosome-bound mRNAs from brain-resident immune cells and subjected them to RNA-seq. We detected high levels of *Wasf1* transcripts and other WRC subunits, including *Cyfip1/2*, *Nckap1*, *Abi1/2/3*, and *Brk1*, in adult hippocampal microglia (Figure 4B). qPCR of TRAP samples suggests robust *Wasf1* translation among *Wasf* paralogs and higher *Abi1* translation compared to *Abi3* or *Abi2* (Figure 4C). Immunohistochemistry revealed WAVE1 protein expression in approximately 70% of microglia in non-Tg mice, with a ∼50% reduction in WAVE1-expressing microglia in P301S tau mice compared to non-Tg mice (Figure 4, E and F). snRNA-seq data further indicate distinct expression patterns of WRC subunit paralogs in non-Tg versus P301S tau mice: *Wasf1* (along with *Wasf3*, *Cyfip2*, *Abi2*, and *Nckap1*) is reduced, *Abi3* (along with *Abi1*, *Nckap1l*, and *Brk1*) is increased, and *Wasf2* and *Cyfip1* remain unchanged in P301S tau mice compared to non-Tg controls (Figure 4, G and H).

**Figure 4.**
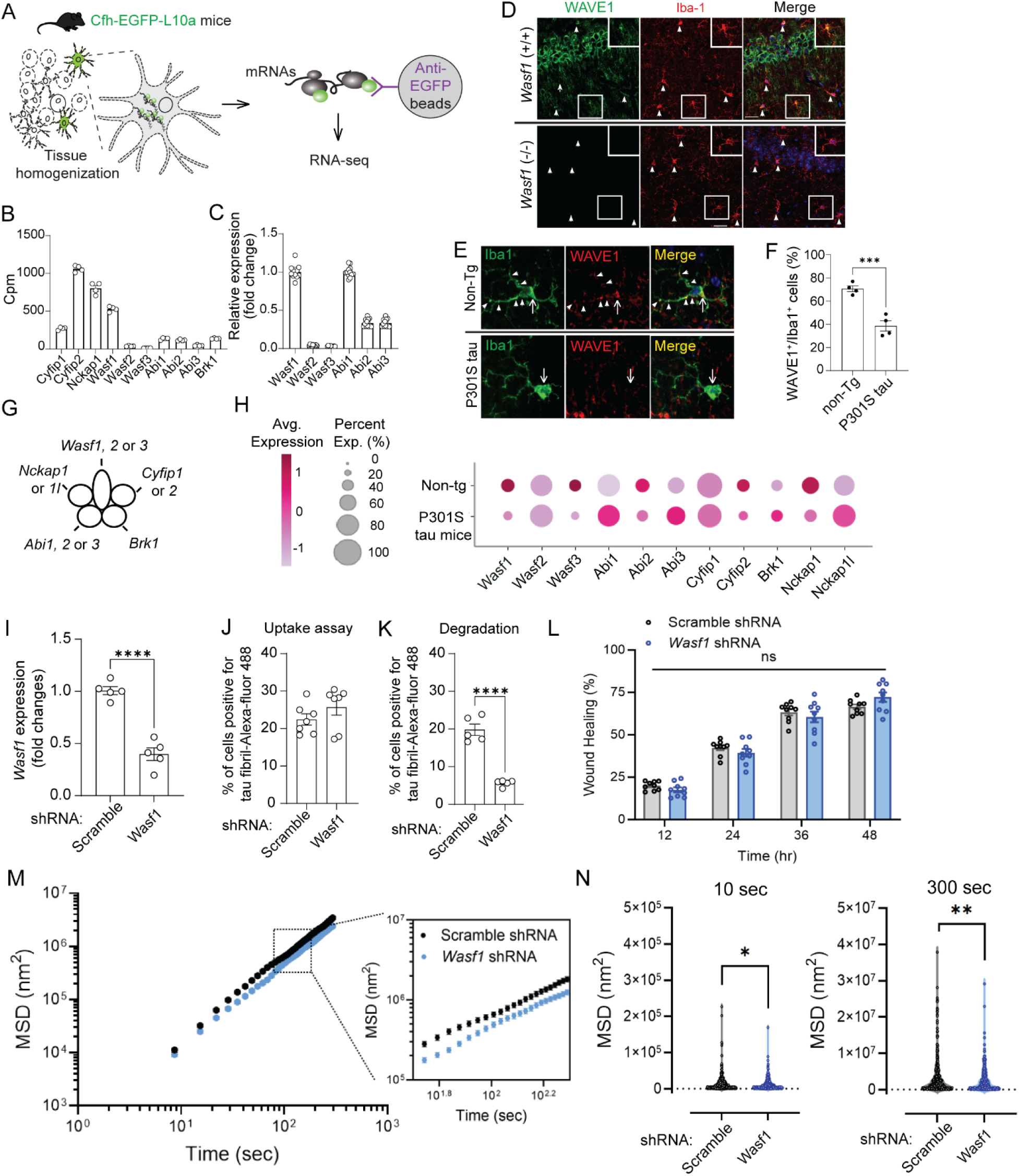
WAVE1 is an endogenous regulator of microglia. (**A**) Schematic of brain immune cell-specific TRAP/RNA-seq using Cfh-EGFP-L10a mice expressing EGFP-fused L10a under the *Cfh* promoter. (**B**) Sequencing-based quantification of mRNAs encoding WRC subunits in 2-month-old Cfh-EGFP-L10a mice (n = 4). (**C**) qPCR of TRAP samples showing expression levels of WAVE and ABI paralogs. (**D**) WAVE1 expression (green) in Iba1-positive cells (red) in the hippocampal CA1 region. WAVE1 −/− panel confirms antibody specificity. Scale bar, 100 μm. (**E** and **F**) Reduced proportion of WAVE1-expressing microglia in the hippocampus of P301S tau mice compared to non-Tg mice. Representative images of Iba1-positive cells (green) with or without WAVE1 expression (red) (**E**). Scale bar: 10 μm. Quantification of WAVE1-expressing microglia (**F**). n = 4 mice/group. Data are mean ± SEM. *\*\*\*P*<0.001, t*-*test. (**G**) Illustration of WRC composition and subunit paralogs. (**H**) snRNA-seq data showing distinct expression patterns of WRC subunit paralogs in non-Tg and P301S tau mice. (**I**) BV2 cells infected with lentiviral vectors expressing scrambled shRNA or WAVE1 shRNA. qPCR was performed to assess knockdown efficiency. Data are mean ± SEM. \*\*\*\**p* < 0.0001, t-test. (**J, K**) Phagocytosis assays using WAVE1-knockdown BV2 cells. Cells were incubated with tau fibril–Alexa 488 for 24 h to measure uptake (**J**). For degradation assays, cells were incubated with tau fibril–Alexa 488 for 24 h, washed, and further incubated for 6 h (**K**). Quantification was performed across 350–900 cells per coverslip (uptake assay: n = 7 coverslips/group; degradation assay: n = 5 coverslips/group). Data are mean ± SEM. \*\*\*\**p* < 0.0001, one-way ANOVA with Tukey’s *post hoc* test. (**L**) Wound closure expressed as a percentage of the initial scratch area at 12, 24, 36, and 48 hours. Data are presented as mean ± SEM. n = 9 technical replicates/group. Two-way ANOVA followed by Šídák’s multiple comparison test. (**M**) Cytoskeletal remodeling of BV2 cells determined by spontaneous nanoscale tracer motion (SNTM) and mean square displacement (MSD) as a function of time. Data are mean ± SEM. n = 304-361 individual bead measurements/group. (**N**) MSD at 10 sec (left) and 300 sec (right). Data are mean ± SEM. \**p* < 0.05, \*\**p* < 0.005, unpaired t-test with Welch’s correction.

To investigate the in-cell effects of *Wasf1* deletion, we first performed phagocytosis assays using a mouse microglial cell line (BV2 cells) and fluorescence-labeled human tau fibrils. BV2 cells were transduced with lentiviral vectors expressing either scrambled shRNA (control) or *Wasf1* shRNA to knock down *Wasf1*. qPCR confirmed ∼60% reduction in *Wasf1* expression (Figure 4I). *Wasf1* knockdown did not alter the ability of BV2 cells to engulf tau fibrils but significantly enhanced the degradation of engulfed tau fibrils (Figure 4, J and K), consistent with increased phagocytic activity previously reported in WAVE1-deficient macrophages (21). On the one hand, *Wasf1* knockdown significantly decreased the rate of cytoskeletal remodeling in BV2 cells (Figure 4, M and N). On the other, local cellular migration (Figure 4L and Supplementary Figure 3, A and B) and the stiffness of BV2 cells (Supplementary Figure 3C) were comparable between *Wasf1* knockdown and control cells. These findings suggest that WAVE1 acts as an endogenous regulator of microglial function and support the in-cell effects of *Wasf1* deletion on microglial activity.

### Wasf1 KO affects microglial interactions with other cells in P301S tau mice

To investigate if *Wasf1* KO alters cellular interactions between microglia and other cell types, we performed the CellChat that infers intercellular communication at the population level by integrating communication probability or strength between sender and receiver cells (visualized as circle or heatmap plots) with single-cell expression heterogeneity data (violin plots) (21). (Figure 5). Information flow analysis, which summarizes overall communication probability among all cell-type pairs in the inferred network, revealed that CD200, Galectin, Activin, TGFβ, CSF, CX3C, and CCL signaling pathways are altered by *Wasf1* KO compared to WT in P301S tau mice (Figure 5A). These signaling pathways are critical for maintaining microglial homeostasis and regulating state transitions under disease conditions (30–34).

**Figure 5.**
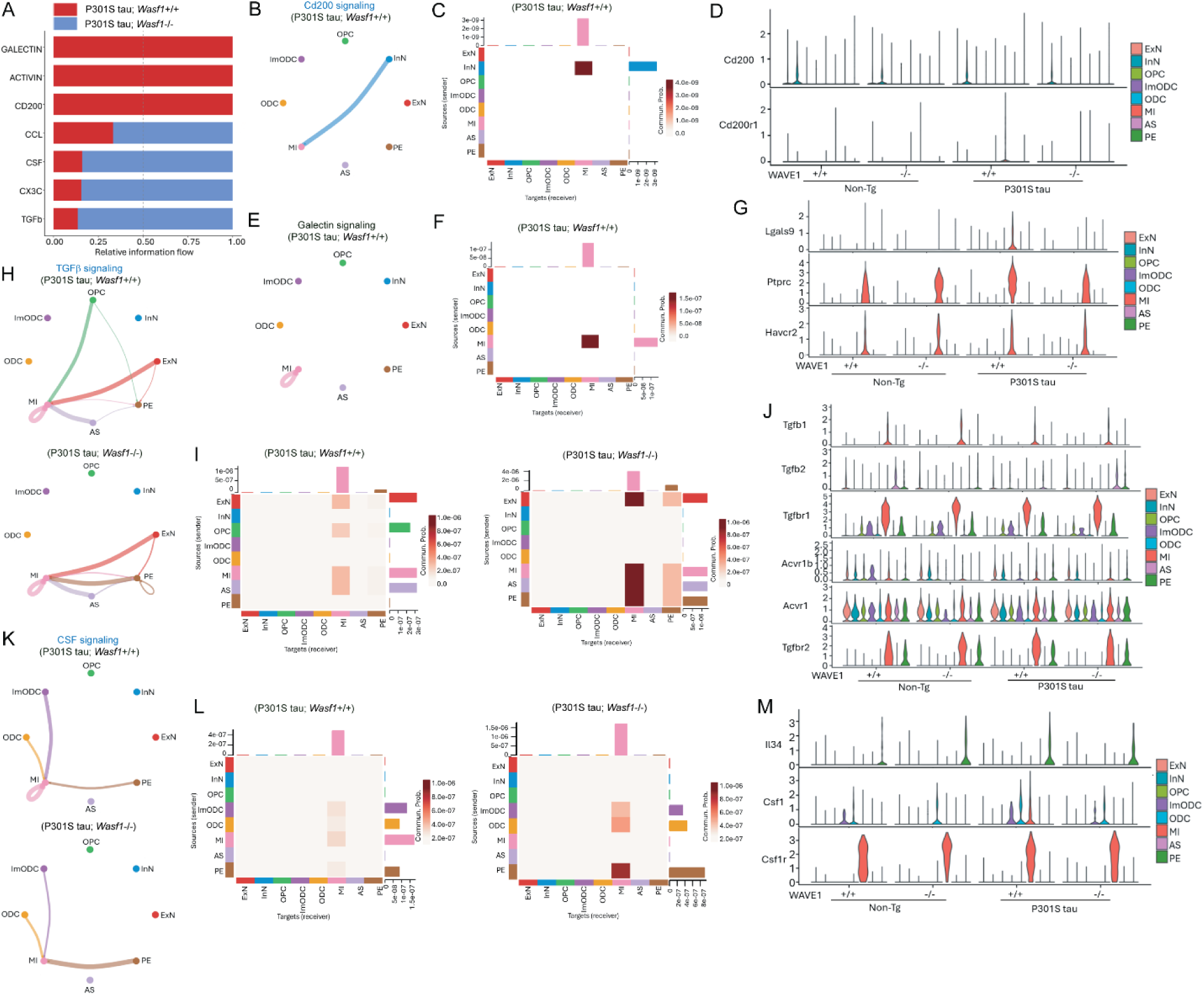
*Wasf1* KO modifies autocrine and cell-to-cell signaling in microglia. (**A**) Information flow chart summarizing overall communication probability among all cell-type pairs in the inferred microglia-related network. (**B**-**D**) CD200 signaling, (**E**-**G**) Galectin signaling, (**H**-**J**) TGFβ signaling and (**K**-**M**) CSF signaling. Circle plots show autocrine and intercellular signaling between cell types in P301S tau mice with *Wasf1*+/+ or −/−. (**B, E, H** and **K**). Heatmaps display communication probabilities aggregated from ligand–receptor interactions for each signaling pathway between cell types in P301S tau mice with *Wasf1*+/+ or −/− (**C, F, I** and **L**). Violin plots illustrate ligand and receptor gene expression comparisons across specific cell types among the four experimental groups (**D, G, J** and **M**).

CD200 is a type I transmembrane glycoprotein, and its receptors are critical for maintaining microglial quiescence and immune homeostasis in the brain (31). CD200 is primarily expressed on neurons, including inhibitory neurons (InN), whereas CD200 receptors are predominantly expressed on microglia (MI). Consistent with the known role of the CD200 pathway in microglial polarization, CD200 signaling is increased in P301S tau mice with WT *Wasf1*. However, *Wasf1* KO reduces this suppressive signaling by lowering CD200 receptor expression to levels comparable to non-Tg (*Wasf1* WT) mice, suggesting restoration of microglial activation by *Wasf1* KO in P301S tau mice (Figure 5, B-D). Previous studies indicate that CD200-deficient microglia exhibit increased phagocytosis associated with enhanced lysosomal activity (35), suggesting a potential link between reduced CD200 signaling and the beneficial effects of *Wasf1* deletion. Galectins, a family of endogenous lectins, function by interacting with numerous glycosylated receptors on immune cell surfaces (36). Galectin-9 expression is significantly elevated in pro-inflammatory and neurodegenerative conditions (37–39). Similarly, Galectin-9 expression is increased in P301S tau mice, but *Wasf1* KO reduces Galectin-9-mediated autocrine activation of microglia in P301S tau mice (Figure 5, E-G). Although violin plots of gene expression for Activin signaling appear comparable between *Wasf1* WT and KO groups in P301S tau mice (Supplementary Figure 4C), the calculated communication probability suggests activation of Activin signaling in *Wasf1* WT but not KO conditions in P301S tau mice (Figure 5A; Supplementary Figure 4, A and B).

Microglia are major recipients of TGFβ signaling originating from excitatory neurons (ExN), oligodendrocyte precursor cells (OPC), astrocytes (AS), as well as through microglial autocrine regulation in P301S tau mice (Figure 5, H-J). However, TGFβ signaling between microglia (MI) and OPC decreased, while MI communication with vascular pericytes (PE) increased (Figure 5H). Overall, communication probabilities among different cell groups were elevated by *Wasf1* KO (Figure 5I). Microglia in P301S tau mice also receive CSF ligands via autocrine signaling and from immature oligodendrocytes (ImODC), oligodendrocytes (ODC), and PE (Figure 5, K-M), as well as CX3C ligands released from ExN and InN (Supplementary Figure 4, D and E). Notably, the signaling flow was significantly altered by *Wasf1* KO compared to WT *Wasf1*. *Wasf1* KO also increased information flow in major chemokine CCL signaling, where microglia act as a minor participant, while ExN and InN serve as major senders, and PE is the primary receiver (Supplementary Figure 4, F-H). Altogether, these results suggest that tau pathogenesis and microglial transitions in P301S tau mice as well as their modulation by *Wasf1* KO are significantly influenced by autocrine and intercellular signaling.

## Discussion

In this study, we identified a pivotal role for WAVE1 in tau pathogenesis and microglial state transitions using the P301S tauopathy mouse model. *Wasf1* KO significantly reduces tau hyperphosphorylation and improves cognition in P301S tau mice. snRNA-seq data indicate that *Wasf1* deletion reverses microglial state transitions and disease-associated microglia (DAM) gene signatures. Although *Wasf1* transcript levels are very low, *Wasf1* mRNA is highly translated in microglia. Furthermore, *Wasf1* knockdown in BV2 cells enhances degradation of engulfed tau fibrils, suggesting a cell-autonomous mechanism underlying the beneficial effects of *Wasf1* deletion. CellChat analysis further demonstrates that *Wasf1* deletion alters microglial autoregulation and their interactions with other cell types, including excitatory and inhibitory neurons, oligodendrocytes, vascular pericytes, and astrocytes, in P301S tau mice. Collectively, these findings suggest that *Wasf1* deletion promotes beneficial microglial phenotypic transitions and mitigates tau pathology, highlighting WAVE1 as a promising therapeutic target for Alzheimer’s disease and related dementias.

Our previous study identified WAVE1 as a key regulator of Aβ production (15). WAVE1 colocalizes and interacts with APP, and WAVE1-driven actin polymerization facilitates vesicle budding and APP trafficking to the cell surface, critical steps for Aβ generation (15). Importantly, we demonstrated consistent downregulation of *Wasf1* gene expression in both human AD brains and AD mouse models (15). Mechanistically, the APP intracellular domain (AICD), generated via the amyloidogenic pathway, binds to the WASF1 promoter and suppresses its transcription and protein expression (15). Because *Wasf1* KO significantly lowers brain Aβ levels and rescues memory deficits in APP/PS1 mice, we proposed a role for WAVE1 in negative feedback regulation of the amyloidogenic pathway (15). Intriguingly, the current study suggests a compensatory role for WAVE1 in tauopathy. Cdk5-mediated inhibitory phosphorylation of WAVE1 is markedly increased in brain lysates, while microglial WAVE1 expression is significantly downregulated in P301S tau mice compared to non-Tg controls, indicating WAVE1 suppression under the tauopathy condition. To investigate the consequences of WAVE1 suppression, we generated a *Wasf1* KO model in P301S tau mice. *Wasf1* KO decreases tau phosphorylation, ameliorates memory deficits, reduces gliosis, and reverses DAM gene expression patterns and microglial state transitions. Thus, the suppression of WAVE1 observed in P301S tau mice appears to represent a cellular compensatory response to tau aggregates. Since WAVE1 is a cellular activity-dependent regulator of actin polymerization (6, 7, 9, 12), elucidating the mechanisms underlying WAVE1 hyperphosphorylation in brain lysates or the downregulation of its gene expression in microglia of P301S tau mice will be an important area of future study.

Previous and current studies have shown the WRC as a critical regulator of microglial function. ABI3 appears to be essential for phagocytosis and neuroprotective microglial activity, as its knockdown or KO impairs phagocytosis, induces a pro-inflammatory phenotype, and facilitates Aβ and tau pathology (18, 20). In contrast, WAVE1 suppresses phagocytosis in macrophages (21) and the degradation of engulfed tau fibrils in microglia, whereas its reduction enhances phagocytic capacity (Figure 4K). Since *Wasf1* KO ameliorates Aβ accumulation in APP/PS1 mice (15), it is possible that *Wasf1* KO facilitates microglia-mediated Aβ clearance. Although impaired migration was reported in *ABI3*-knockdown microglial cells (18), we observed no change in migration following *Wasf1* knockdown in BV2 cells, despite a significant reduction in actin cytoskeleton dynamics (Figure 4, L–N; Supplementary Figure 3, A and B). Other WRC subunits have also been implicated in microglial function. WAVE2 enhances lipopolysaccharide (LPS)-induced microglial phagocytosis when overexpressed (40) and interacts with LRRK2 in Parkinson’s disease models (40, 41). *CYFIP1*, a gene associated with autism spectrum disorders and schizophrenia, is expressed in microglia and regulates synaptic pruning and neurogenesis (42, 43). *NCKAP1* reduction is associated with defective phagocytosis and exaggerated inflammation in ALS patient-derived microglia-like cells, whereas *NCKAP1* overexpression rescues these phenotypes (44). Collectively, WRC subunit paralogs exhibit distinct, and in some cases opposing, roles in microglial function and state transitions, although the mechanisms by which varying levels of these paralogs fine-tune WRC-mediated actin polymerization dynamics and microglia functional regulation remain largely unknown.

A previous study reported that microglial activation is an early event in P301S tau mice, and administration of FK506, a potent immunosuppressive drug, reduces tau pathology (24). Another study demonstrated that experimental inhibition of NF-κB signaling in microglia rescues learning and memory deficits in P301S tau mice, establishing a causal link between aberrant microglial activation, tau pathology, and cognitive impairment (25). Consistent with these findings, our snRNA-seq data indicate that gene expression changes in microglia are most pronounced in P301S tau mice compared to non-Tg controls or *Wasf1* KO versus *Wasf1* WT conditions in P301S tau mice. In contrast, the effect of *Wasf1* KO on gene expression in other cell types was modest in P301S tau mice (Supplementary Figure 2). Interestingly, *Wasf1* KO induces extensive gene expression changes in inhibitory neurons, oligodendrocytes, and astrocytes, but not in microglia, in non-Tg mice (Supplementary Table 1). Notably, our CellChat analysis indicates that the effects of *Wasf1* KO on microglial state changes are significantly influenced by both autocrine and intercellular signaling mechanisms. Therefore, we cannot rule out the possibility that the beneficial effects of *Wasf1* deletion in P301S tau mice were driven, at least in part, by *Wasf1*-deleted neurons or other non-microglial cell types. Future studies should employ microglia-specific or other cell-type-specific KO approaches to delineate the cell-type-specific roles of WAVE1 in AD pathogenesis. Despite this limitation, our study reveals for the first time a critical role for WAVE1 in microglial function and suggests that distinct WRC subunits may serve as potential therapeutic targets for modifying maladaptive microglial responses in tauopathy-related dementias and other neurodegenerative diseases.

## Methods

### Sex as a biological variable

Because sex differences in tau pathogenesis in Alzheimer’s disease and related dementias are well documented(45, 46), we analyzed males and females as separate groups in most experiments. For snRNA-seq experiments, however, we combined data from both sexes.

### Mice

P301S tau (PS19 model) was purchased from the Jackson Laboratory (Strain #:008169). The constitutive *Wasf1* KO mice were generated as previously described (15). We produced the progeny of P301S tau mic crossed with *Wasf1* KO mice by natural breeding or *in vitro* fertilization (IVF) and embryo transfer techniques (Genome Editing Shared Resource, Rutgers University) to provide animals for experiments.

All mice are of C57BL/6 background. Mice were housed 2-5 per cage with a 12-hour light/12-hour dark cycle and *ad libitum* access to food and water under controlled humidity, temperature (21±2 °C), and air exchange. Mice were assigned to experimental groups based on their genotype. The selection of animal samples from different experimental groups for biochemical analyses was performed randomly and blindly.

### Immunoblotting

Brains were dissected on ice, and hippocampi were snap-frozen in liquid nitrogen and stored at -80°C until further experiments. Dissected hippocampi samples were lysed with RIPA buffer (#R0278, Sigma-Aldrich), protease inhibitor cocktail tablets (#04693124001, Roche), and phosphatase inhibitor cocktail tablets (#04609837001, Roche) and homogenized with a sonicator (#CL-188, QSonica) for 10 sec twice at a cold room. Then, the samples were centrifuged at 15,000 × g for 30 min at 4°C. Small aliquots were reserved for protein determination using a BCA protein assay kit (#23225, Thermo Scientific) with bovine serum albumin as the standard.

The remaining lysates were mixed with sample buffer containing 50 mM Tris-glycine (pH 6.8), 2 % sodium dodecyl sulphate (SDS), 100 mM dithiothreitol (DTT), 10 % glycerol and 0.05 % bromophenol blue. The mixtures were boiled for 5 minutes at 95°C. Equal amounts of protein per sample (10 µg) and pre-stained protein size markers (#161-0394, Bio-Rad) were separated by SDS-PAGE 4–20%, Tris-glycine gels (#XP04205BOX, Invitrogen), and separated proteins were transferred to nitrocellulose membranes (#1620233, Bio-Rad). Membranes were blocked with TBS-T buffer (Tris-buffered saline with 0.1% Tween 20) containing 5% non-fat milk. The blot was incubated overnight. at 4°C with the following primary antibodies: WAVE1 (C-terminus, 1:500)(9), WAVE1 p-S310 (1:500)(9), WAVE1 p-S397 (1:500)(9), WAVE1 p-S441 (1:500)(9), tau (tau 46, #4019S, 1:500, Cell Signaling), p-tau S199 (#29957S, 1:500, Cell Signaling), p-tau T231 (#71429S, 1:500, Cell Signaling), p-tau Ser202-Thr205 (AT8, #MN1020, 1:500, Invitrogen), or Gapdh (#MAB374, 1:1000, Millipore). The membranes were washed for 30 min with TBS-T and then horseradish peroxidase (HRP)–conjugated goat anti-rabbit or HRP-conjugated goat anti-mouse IgG secondary antibodies (1:5000, ThermoFisher Scientific) were used at 1:5000 for 1 hour at room temperature (RT). The chemiluminescence signals were developed using Clarity Western ECL substrate (#170-5060, BioRad). The images were acquired in a ChemiDoc Imaging System 3.0.1.14 (Bio-Rad), and the densitometry of the bands was quantified using FIJI software.

### Immunohistochemistry

Mice under CO_2_ anesthesia were transcardially perfused with 10 ml of cold PBS followed by 40 ml of 4% cold paraformaldehyde (PFA) in 0.1 M PBS (pH=7.4). Brains were removed from the skull and postfixed for 6 hour in PFA followed by 24 hour in 30% sucrose at 4°C. 20-µm coronal sections were cut in a cryostat, mounted on glass slides, and stored at -80°C until use.

Brain slides were thawed and dried for 1 h at RT and washed three times with 10 mM PBS. Then, the brain sections were blocked with a solution containing 0.1% Triton X-100 (#X100-100ML, Sigma-Aldrich) and 10% normal goat serum (#S-1000, Vector Laboratories) at RT for 1 hour and incubated overnight at 4°C with one of the following primary antibodies: anti-Iba1 (chicken polyclonal, 1:250, #PA5-143572, Invitrogen) or anti-WAVE1 (rabbit polyclonal, 1:500, V0101 (6, 47)). Then, sections were washed three times with PBS and incubated with secondary antibodies: goat anti-chicken conjugated with Alexa-fluor 488 (1:500, #A11039, Invitrogen) or donkey anti-rabbit conjugated with Alexa-fluor 555 (1:500, #A31572, Invitrogen) at RT for 1 hour, respectively. Then, sections were washed three times with PBS, and slides were mounted with Vectashield Vibrance Antifade Mounting Medium with DAPI (#H-1800-10, Vector Lab), dried overnight, and stored at 4°C until image acquisition. Negative controls were made by omitting the primary antibody.

The hippocampal area (CA1, CA2, CA3, and DG) from bregma -1.70 and -1.82 mm was observed with an epifluorescence microscope with the 20X non-oil immersion lens (DM6R, Leica) and the software Leica Application Suite X (LAS X, Leica). A confocal microscope (Dragonfly 200, Andor Technology, Oxford Instruments) equipped with a Zyla sCMOS camera, and with the 20X non-oil immersion and 40X oil immersion lens, and a 2X digital zoom with the software Fusion (Oxford Instruments). DAPI was excited using a 405 nm emitting diode laser, Alexa Fluor 488 was excited using an argon laser (488 nm), and Alexa Fluor 555 was excited using a 532 nm DPSS laser. The obtained images were analyzed with FIJI software.

### Behavioral tests

Behavioral experiments were conducted between 9:00 a.m. and 5:00 p.m. The experimenters were blind to all genotypes. All tests were carried out using a video-tracking system EthoVision-XT 14 video tracking (Noldus Information Technology, Leesburg, VA, USA). All the apparatus were cleaned with 70% ethanol and dried between each mouse to eliminate the effects of odor. At the end of the experimental testing, mice were returned to their home cage and transferred back to colony housing.

Open Field Test (OFT): To assess mouse locomotion, the mice were individually placed in the center of the OF apparatus (a square arena of 40 cm × 40 cm enclosed by continuous 50 cm high walls) and allowed to freely explore it for 10 min while recording with a camera. Locomotor behavior was examined by determining the total distance travelled and the total time spent in the center or the periphery.

Y-maze Test (YMT): The YMT assesses working memory based on the innate preference of a mouse to alternate arms while exploring a new environment in the apparatus, which consists of three arms: 25 cm (length) × 8 cm (height) × 5 cm (width). Typically, mice prefer to explore a new arm of the maze rather than returning to the one that was previously explored. Each mouse was placed at the end of one arm and allowed to explore the maze freely for 6 minutes. With a camera, the sequence and total number of arms entered were recorded. An entry into an arm was considered valid when all four paws entered the arm. An alternation was defined as entries in different arms. The percentage alternation score and the total distance moved were calculated.

Morris Water Maze (MWM): MWM assesses hippocampal-dependent spatial learning and memory in mice. Testing occurred in a white circular tank (120 cm diameter and 90 cm deep; ENV594M-B, Med-Associates) filled with water (24 ± 1°C) made opaque with white, non-toxic tempera paint. The tank contained a transparent acrylic escape platform (10.2 cm in diameter) in the northeast quadrant of the tank, which remained there for all five days of the test. Visible spatial cues of equal size, surrounded the tank, facing each other, were placed at North (N; star), South (S; circle), East (E; square), and West (W; triangle) coordinate locations within the tank, which divided the tank into NE, SE, SW, and NW quadrants. Data was collected using a video tracking system. A four-trial procedure was conducted over four consecutive days, preceding the testing day (day five).

In the first trial day, the platform was placed visible (0.5 cm outside the water) with an orange flagpole protruding 8 cm above the water’s surface so the mice could see it. In the following days, the platform was underwater (1 cm covered with water), and the pennant was removed. To measure learning during MWM training, mice were released from NE, SE, SW, NW points where swim latency (s) to escape onto the hidden platform were video tracked, traced, and analyzed. If the mouse did not find the platform within 1 min, then it was led to the platform by the experimenter. Upon reach or located on the platform it sat for 15 s before being removed. The mouse was then dried and returned to the home cage. The latency to the platform (s) was recorded for each trial.

On the fifth day, the day of the test, the platform was removed from the tank, and the mouse was placed in the tank in the quadrant opposite the platform for 1 min. The amount of time the mouse spent swimming in the target quadrant (the quadrant that contained the platform during the trial sessions) and the total distance moved (in centimeters) were recorded.

### BV2 cell cultures and Wasf1 knockdown

Mouse microglial BV2 cells were cultivated in DMEM (#11995-065, Gibco) containing 10% fetal bovine serum (FBS) (#A56695, Gibco), and 1% antibiotic-antimycotic (#15240-062, Gibco) at 37°C and 5% CO_2_. The medium was changed three times a week, and cells were maintained at ∼50-70% confluency to avoid activation from overcrowding.

2×10^5^ BV2 cells/ml were seeded in a 12-well plate. After 24 hours, the media was replaced with DMEM containing 7 μg/ml polybrene (#TR-1003-G, Millipore) and cells were incubated with the lentivirus particles (LVRU6P, GeneCopoeia) expressing shRNA scramble control or WAVE1 shRNA (MOI=10). After 48 h, the cells were transferred to a 6-well plate, and the media was replaced with fresh complete media. After 48 hours, the media was changed and supplemented with 3 μg/ml of puromycin (#A11138-02, Gibco). Over consecutive days, the media was changed daily, increasing the concentration of puromycin to 10 μg/ml. Each cell line was maintained in a separate T75 flask, and the media were always supplemented with 10 μg/ml of puromycin. Knockdown was confirmed by qPCR.

### Phagocytosis assays

Pre-formed fibrils of human tau-441 K18 (P301L) (TAU-H5113, Acro Biosystems) were used to conjugate with Alexa Fluor 488 as described (48). Briefly, tau protein was incubated with BRB80 buffer (80 mM PIPES, 1 mM MgCl_2_, and 1mM EGTA, pH 6.8) with a 10-fold molar excess of tris-(2-carboxyethyl) phosphine at room temperature for 30 min. Then, a 4-fold molar excess of Alexa Fluor 488 C-5 maleimide (A10254, Invitrogen) dissolved in DMSO was added and incubated at RT for 3 h in the dark following the manufacturer’s instructions. The unlabeled fluorophores were separated from the labeled protein solution using a NAP-5 column (17085301, Cytiva) following the manufacturer’s instructions. The protein concentration was determined by BCA, and the conjugated tau aliquots were stored at 80°C until use.

For the tau phagocytosis assay (25, 49), BV2 cells were plated in 24-well plates with glass coverslips for 24 hours. Then, BV2 cells were incubated with tau fibrils conjugated with Alexa Fluor 488 (2.5 µg/ml) for 24 hours. To measure tau uptake, BV2 cells treated for 24 hours with tau were incubated with DMEM containing 0.01% trypsin for 5 minutes to remove tau from the cell surface. After incubation, the cells were washed and fixed with cold methanol for 10 minutes. To measure tau degradation, after a 24-hour incubation with tau fibrils, cells were incubated with DMEM containing 0.01% trypsin for 5 minutes, and complete DMEM media was added for 6 hours to allow for tau degradation. After, cells were washed and fixed with cold methanol for 10 minutes. Cell images were captured using a confocal microscope (Dragonfly 200, Andor Technology) at 20X and 40X magnification. The number of tau-positive cells was counted using the multi-point tool in FIJI software.

### TRAP, RNA isolation, reverse transcription, and Quantitative real-time PCR (qPCR)

TRAP was conducted as previously described (29). Total RNA was extracted using the RNeasy Plus Micro Kit (#74034, Qiagen) according to the manufacturer’s protocol. One microgram of total RNA was converted to cDNA using GoScript Reverse Transcription Mix kit (#A2791, Promega) according to the manufacturer’s instructions. qPCR was performed with the PowerUp SYBR Green Master Mix (#A25742, Applied Biosystems). The mouse *Gapdh* primers were F 5’- CAT CAT TGC CAC CCA GAA GAC TG -3’ and R 5’- ATG CCA GTG AGC TTC CCG TTC AG -3’, the mouse *Wasf1* primers were F 5’- CTG CCT GTA ATC AGT GAC GCA AG -3’ and R 5’- GCT TCC TGT TCA CGC TGC TCT T-3’, and the mouse *Abi3* primers were F 5’- GGC AGG TAG AAG CCA AGA TGA G -3’ and R 5’- GGG ATG ACT TTC TGG TTA GAG GG -3’. For the qPCR analysis of RNAs isolated from Cfh-EGFP-L10a TRAP line, the *Wasf1* primers were F 5’-AAG GTA TTC AGC TTC GCA AAG TG-3’ and R 5’-CAA TCC GCT CGT GTT TTG CTT C-3’, the *Wasf2* primers were F 5’-CCA AAT AGA GGG AAT GTG AAC CC-3’ and R 5’-AGG TGG ACG AAT ACC CAG AAG-3’, the *Wasf3* primers were F 5’-AGA TGA CAA GAA GGA TGG GCT G-3’ and R 5’-CCA GAG GTC AAA GAA GTA AGA-3’, the *Abi1* primers were F 5’-GGA AGT AGT GGA GGA AGC GG-3’ and R 5’-GCA CAG CAA TAG GAA TGC CAA TG-3’, the *Abi2* primers were F 5’-GTT GTG GCA ATT TAT GAT TAT ACA AAA G-3’ and R 5’-ATA TAA TGG CTC CTT CCT GAA AGG-3’ and *Abi3* primers were F 5’-CTA CTG CGA GGA TAA CTA CTT GC-3’ and R 5’-CAG GTT ACC CAC TTG GTA GGC-3’.

Reactions were run at 50°C for 2 min and 95°C for 10 min, followed by 40 cycles of 95°C for 15 s and 60°C for 1 min on a Thermal Cycler (#T-100, Bio-Rad). Comparative quantitation of each target gene was performed based on the ΔCT value, which was normalized to the *Gapdh* gene.

### Wound healing migration assay

BV2 cells were seeded at a density of 60,000 cells/well on a tissue culture-treated 96-well plate overnight. Uniform scratches were simultaneously made in each well using a JuLI Stage Scratcher (NanoEntek), and then the wells were washed twice with media. Images were taken every 20 minutes for 48 hours using a MuviCyte Live-Cell Imaging System (PerkinElmer), housed in an incubator maintained at 37°C and 5% CO2. The migration of BV2 cells into the wound over time was analyzed as a percentage of the wound area at t = 0 minutes.

### Spontaneous and forced bead motions with magnetic twisting cytometry (MTC)

For these studies, BV2 cells were seeded at a density of 60,000 cells/well on tissue culture-treated plastic wells in a 96-well format. The next day, cells were incubated for 20 min with RGD-coated ferrimagnetic microbeads (∼4.5 µm in diameter), washed, and spontaneous nanoscale displacement of individual microbeads (∼50 beads per field of view) functionalized to the cytoskeleton through cell surface integrin receptors was first recorded at a frequency of 12 frames/s for t_max_ ∼ 300 s via a CCD camera (Orca II-ER, Hamamatsu) to assess cytoskeletal remodeling dynamics (50, 51). Bead trajectories in two dimensions were characterized by computing mean squared displacement of all beads as a function of time [MSD(t)] (nm^2^) as previously described (50). We then applied forced motions of the functionalized microbeads using MTC as previously described (51) to measure the stiffness of individual cells. In brief, the RGD-coated ferrimagnetic microbeads were magnetized parallel to the cell plating (1,000 Gauss pulse) and twisted in a vertically aligned homogenous magnetic field (20 Gauss) that was varying sinusoidally in time. The sinusoidal twisting magnetic field caused both a rotation and pivoting displacement of the beads that lead to the development of internal stresses that resist its motion. Lateral bead displacement was optically detected with a spatial resolution of 5 nm, and the ratio of specific torque to beads displacement was computed and expressed as the cell stiffness (Pa/nm). The same population of cells (with attached RGD beads) was used to acquire both the cytoskeletal rearrangement and stiffness measurements in the same experiment.

### Nuclei isolation from frozen mouse hippocampus

Nuclei were isolated from one hemisphere of the mouse hippocampus of frozen brain tissue. Briefly, frozen tissue was homogenized on ice in freshly prepared Tissue Homogenization Buffer (THB; 320 mM sucrose, 10 mM Tris-HCl (pH 7.4), 10 mM NaCl, 3 mM MgCl₂, 0.1% NP-40, 1 mM DTT, 50 U/mL DNase I) supplemented with RNase inhibitor. Homogenization was performed using a pre-chilled Dounce homogenizer (10 strokes with the loose pestle and 15 strokes with the tight pestle). The homogenate was carefully layered onto a 1.8 M sucrose cushion (10 mM Tris-HCl (pH 8.0), 3 mM Mg-acetate) and centrifuged at 10,000 × g for 20 min at 4 °C. The supernatant was removed, and the nuclear pellet was gently resuspended in cold wash buffer (1× PBS with 2% BSA and 0.2 U/μL RNase inhibitor). Nuclei were filtered through a 40 μm cell strainer, washed, and pelleted at 500 × g for 5 min at 4 °C. The purified nuclei were subsequently fixed using the Parse Biosciences Nuclei Fixation Kit, following the manufacturer’s instructions. Final nuclei suspensions were counted using a hemocytometer and stored at -80 °C until snRNA-seq library preparation.

### snRNA-seq barcoding and library construction

snRNA-seq libraries were constructed using the Whole Transcriptome v3 kit (Parse Biosciences) following the manufacturer’s protocol. In brief, nuclei derived from different samples were multiplexed and barcoded using the SPLiT-pool combinatorial barcoding method (52). During this process, individual nuclei undergo four sequential rounds of split-pooling across 96-well plates containing well-specific oligonucleotide barcodes, ensuring that each nucleus receives a unique barcode combination. All samples were processed and barcoded simultaneously to minimize batch effects associated with separate library preparations. The resulting libraries were sequenced on an Illumina NovaSeq X Plus platform with paired-end 150 bp reads. Sequencing was targeted at approximately 60,000 reads per nucleus, yielding an average read depth of 64,241 reads per nucleus.

### Data preprocessing and quality control

Demultiplexed FASTQ files from each sample were uploaded and processed using the Trailmaker^TM^ pipeline (Parse Biosciences). The pipeline generated digital nucleus × gene count matrices from the raw sequencing data, yielding a total of 93,702 nuclei prior to quality control (QC). Nuclei QC was performed automatically within the Trailmaker^TM^ Insights module, which applies sample-specific thresholds to filter low-quality nuclei and identify potential doublets. Automatic filtering was based on the distributions of total UMI counts, number of detected genes per nucleus, and mitochondrial RNA read fractions. After QC filtering, 75,948 high-quality nuclei were retained for downstream analyses. Dimensionality reduction and data embedding were performed within the Insights module using the default Trailmaker^TM^ workflow, which incorporates principal component analysis (PCA) and Uniform Manifold Approximation and Projection (UMAP)–based visualization methods. All analyses and visualizations were performed within Trailmaker^TM^.

### CellChat analysis

The CellChat (v2.1.1) (https://github.com/jinworks/CellChat?tab=readme-ov-file) package has been utilized for the inference, analysis, and visualization of cell-cell communication from single-cell transcriptomics data (53). We used snRNA-seq data for CellChat analysis. An individual CellChat object was generated using the createCellChat function, which utilized the normalized data matrix and cell annotation from the Seurat object. Subsequently, ligand-receptor gene pairs, which were overexpressed in each cell type, were identified with a p-value 0.05 cutoff. A permutation test was applied to calculate the significance of cell-cell communication (communication probability) using the computeCommunProb function. The communication probabilities of ligand-receptor gene pairs were summarized in each signaling pathway by computeCommunProbPathway function. Finally, signaling pathways of interest were visualized using circle plot, heatmap, and violin plot.

### Statistics

All data are expressed as means ± SEM except violin plots and cell proportion graphs. Sample sizes for biochemistry, behavioral, and imaging data were determined based on our empirical data accumulated in the laboratory. Sample sizes and statistical methods are provided in each figure legend. GraphPad Prism version 10 was used for statistical analysis and graphical preparations. *p*< 0.05 was considered the cutoff for statistical significance. Statistical significance is shown as \**p*<0.05, \*\**p*<0.01, \*\*\**p*<0.001, \*\*\*\**p*<0.0001, otherwise indicated in figures or figure legends.

### Study approval

All procedures for animal crossings, biochemical, histochemical and behavioral experiments were approved by Rutgers University Institutional Animal Care and Use Committee (PROTO202100001) and were in accordance with the National Institutes of Health guidelines.

### Data availability

snRNA sequencing datasets were deposited in the NCBI’s Gene Expression Omnibus (accession number GSE316281). Raw data related to the figures in this article may be available upon request to the corresponding author.

## Author contributions

CB, W-HC, JHL, Y-SK, SSA, and YK conceptualized and designed the study. CB and YK generated experimental mice. CB, W-HC, JL, JHL, and YYJ performed the experiments. AA and YT performed CellChat analysis. J-HA provided critical resources. CB, W-HC, AA and YK wrote the manuscript. All authors approved the manuscript.

## Acknowledgments

We thank Anandakumar Shunmugavel, Bruno Carabelli, and Sang Hoon Kim for their technical assistance with biochemical and behavioral assays. We also thank Peter J. Romanienko at the Rutgers Genome Editing Shared Resource for providing excellent *in vitro* fertilization and embryo transfer services. This project has been funded by a grant from New Jersey Health Foundation (to YK), a Busch Biomedical Grant from the Office for Research at Rutgers (to YK), an International Collaborative Research Grant from Rutgers Global at Rutgers (to YK), and the Rutgers Startup Fund (to YK). YK was supported in part by NIH R01MH121763 and R21NS130250. SSA was supported by NIH R01HL164404. JHA was supported by the National Research Foundation of Korea (NRF) grant funded by the Korea government (MSIT)(RS-2023-NR076509) and the Ewha Global Excellence Program Research Grant.

---

**Supplementary Figure 1.**
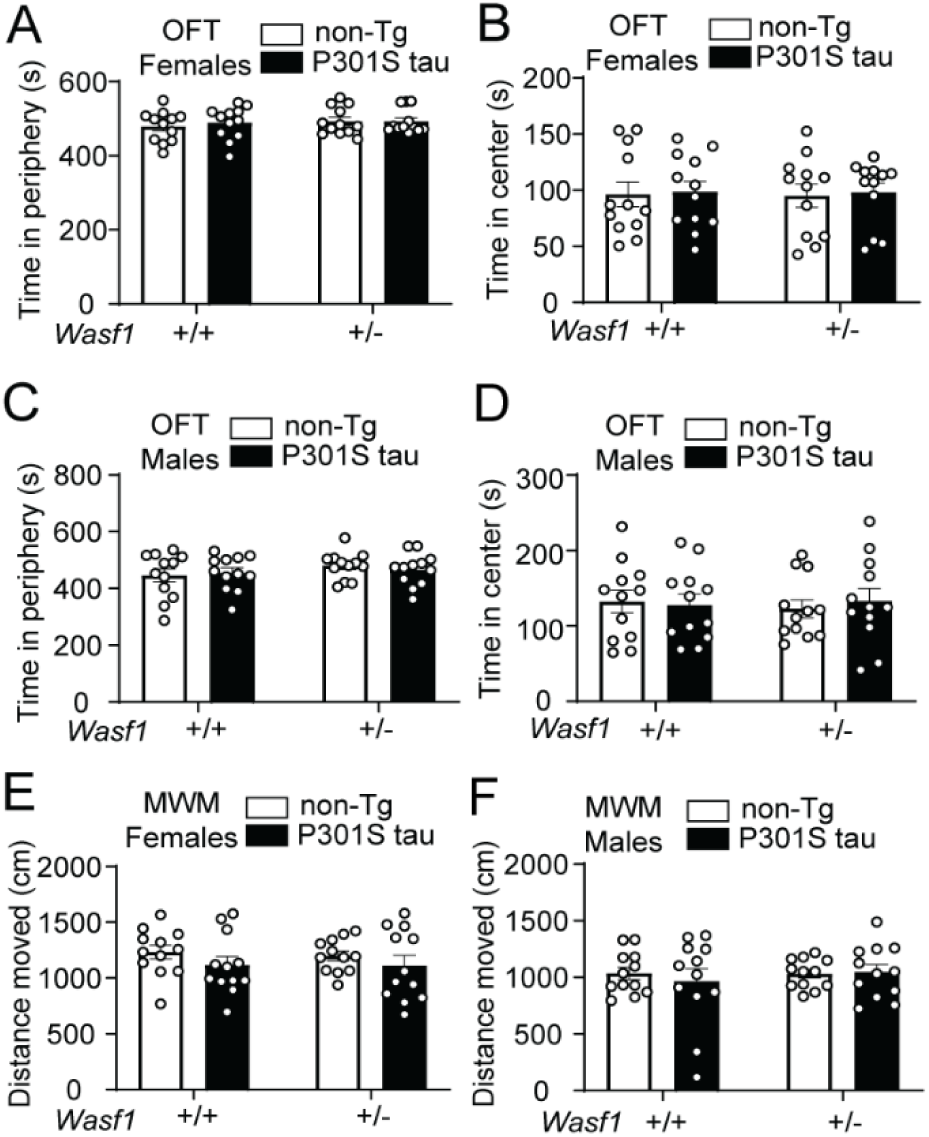
Further analyses of locomotor activity in the open field and Morris water maze tests. (**A** and **B**) Time spent in the periphery (left) and center (right) in the open-field test for females (**A**) and males (**B**). (**C**) Total distance moved by females (left) and males (right) during the probe trial assessing target platform acquisition. Mice aged 6-8 months (n = 12/group). No significant differences were observed between groups. Two-way ANOVA with Tukey’s *post hoc* test.

**Supplementary Figure 2.**
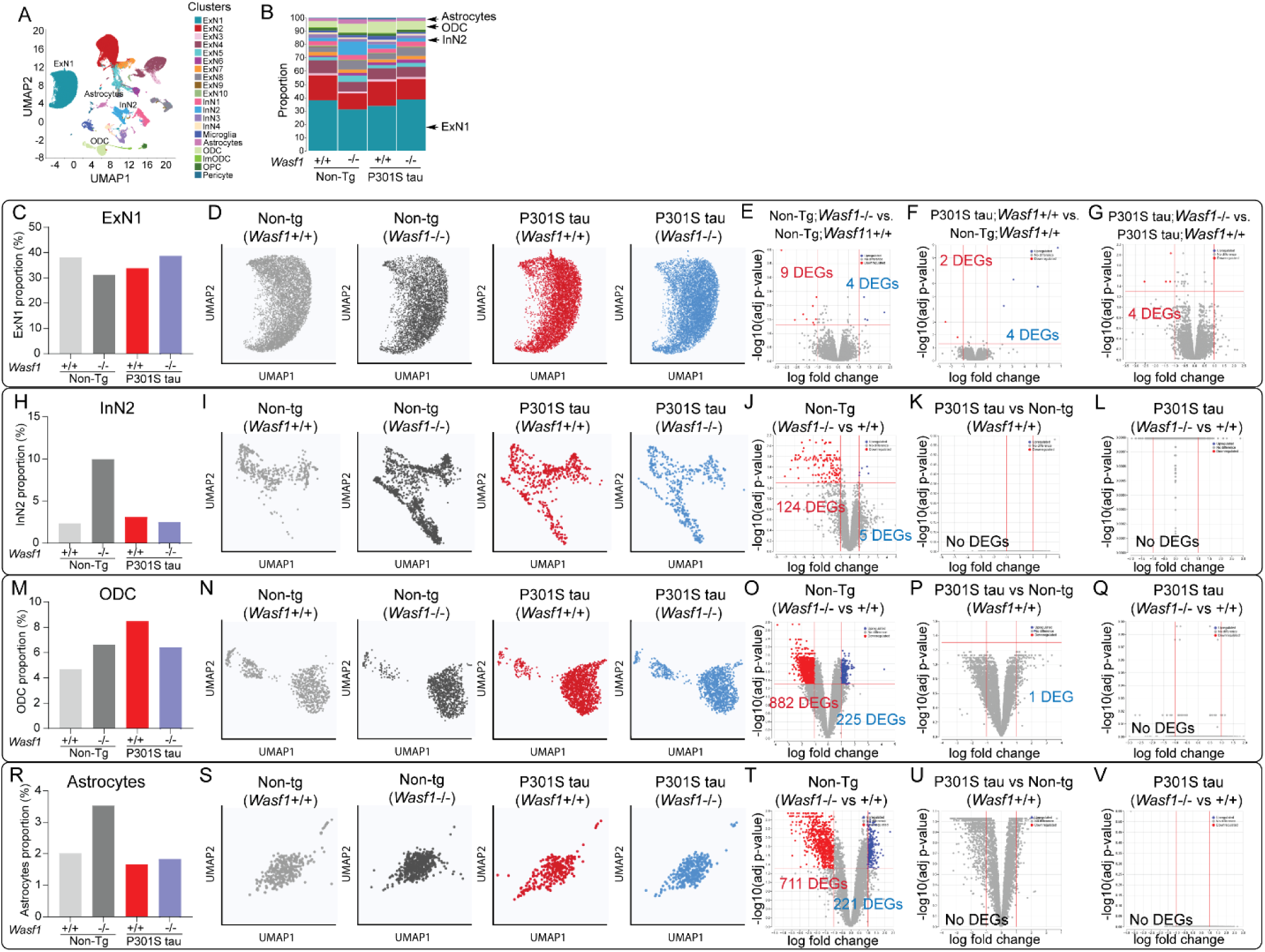
snRNA-seq datasets for other cell types. (**A**) UMAP visualization of all cell clusters. (**B**) Proportion of specific cell types in each group. (**C**-**V**) Detailed analyses for individual cell types. Proportion of specific cell types in each group (**E**, **H**, **M** and **R**). UMAP plots highlighting subpopulations of specific cell types (**D**, **I**, **N** and **S**). Volcano plots showing DEGs in ExN1 (**E**-**G**), InN2 (**J**-**L**), ODC (**O**-**Q**), Astrocyte (**T**-**V**). Mice aged 8-9 months. *n* = 6 per group (3 males and 3 females).

**Supplementary Figure 3.**
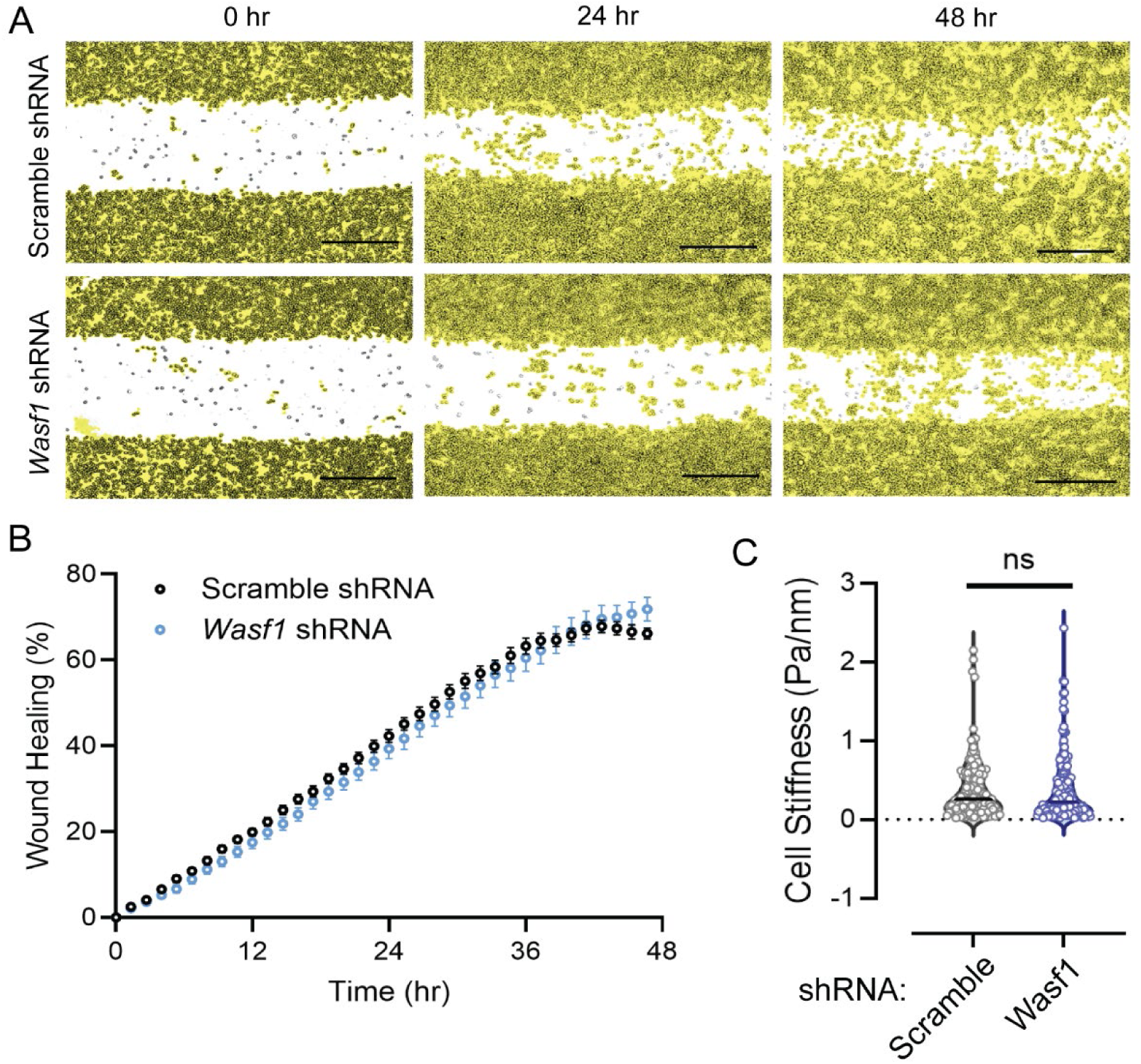
Wound healing and cell stiffness assays using BV2 cells transfected with lentiviral vectors expressing shRNA. (**A**) Representative images of wound healing assays. Scale bars, 500 µm. (**B**) Migration of BV2 cells measured every 20 min for 48 h using wound-healing assay. Data are mean ± SEM. n = 9 technical replicates/group. (**D**) Optical magnetic twisting cytometry (MTC) used to assess cell stiffness.

**Supplementary Figure 4.**
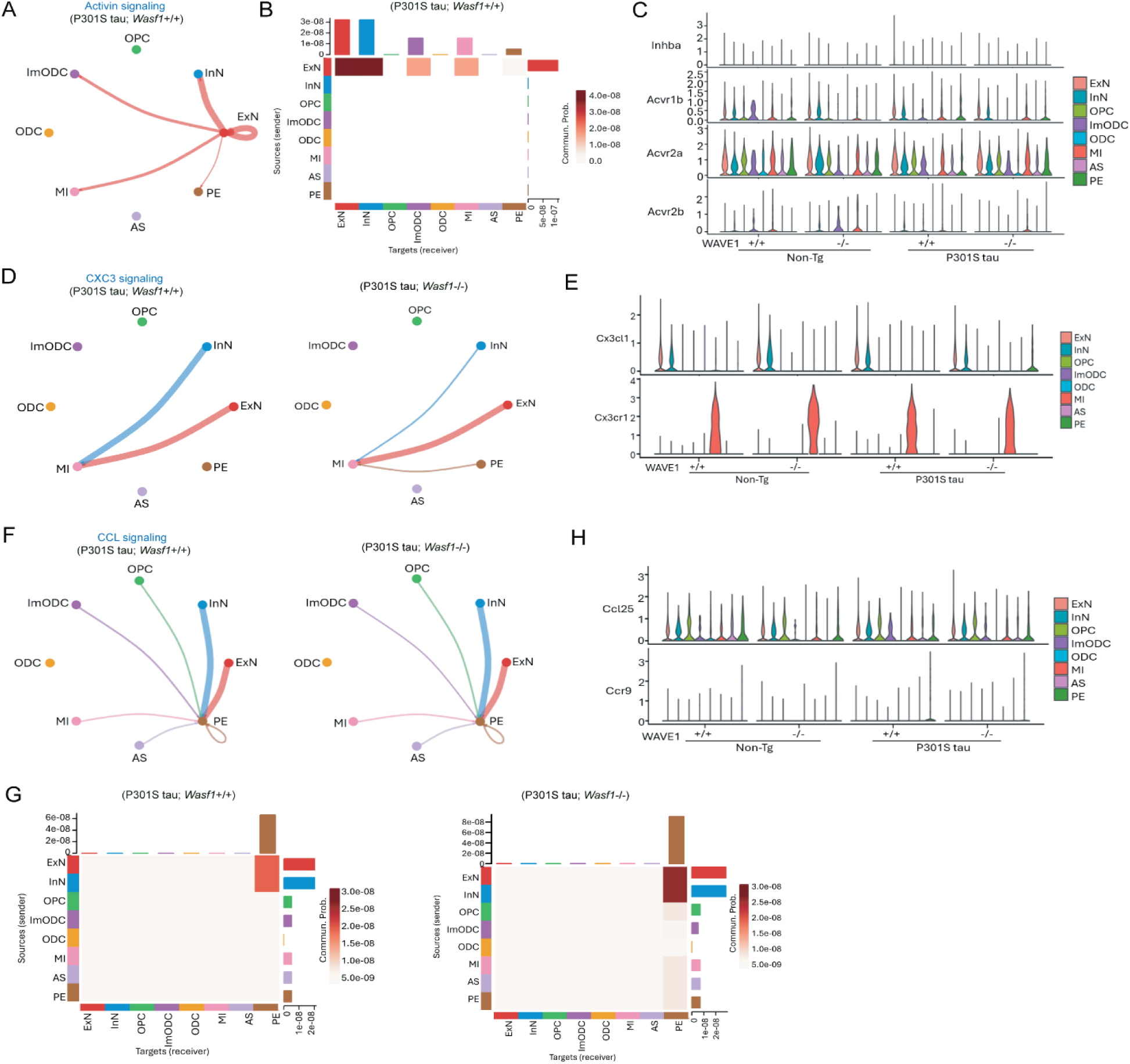
*Wasf1* KO alters Activin, CX3C, and CCL signaling in P301S tau mice. (**A**-**C**) Activin signaling, (**D** and **E)** CXC3 signaling, and (**F**-**G**) CCL signaling. Circle plots showing autocrine and intercellular communication between cell types in P301S tau mice with *Wasf1*+/+ or *Wasf1*1-/- (**A**, **D** and **F**). Heatmaps displaying communication probabilities aggregated from ligand-receptor interactions within each signaling pathway between cell types in P301S tau mice with *Wasf1*+/+ or *Wasf1*1-/- (**B** and **G**). Violin plots comparing ligand and receptor gene expression across specific cell types among the four experimental groups (**C, E** and **H**).

